## Supplementary Figures for "Analysis of genetically independent phenotypes identifies shared genetic factors associated with chronic musculoskeletal pain at different anatomic sites"

### Figure S1.

Graphical summary of the discovery GWAS stage after the genomic control correction using LD Score regression intercept.

Red line corresponds to the genome-wide significance threshold of *P* = 1.25e-08 (5.0e-08/4, where 4 is the number of GIPs). Only associations with
*P* < 1.0e-02 are presented. Replicated loci are annotated.

**GIP1**

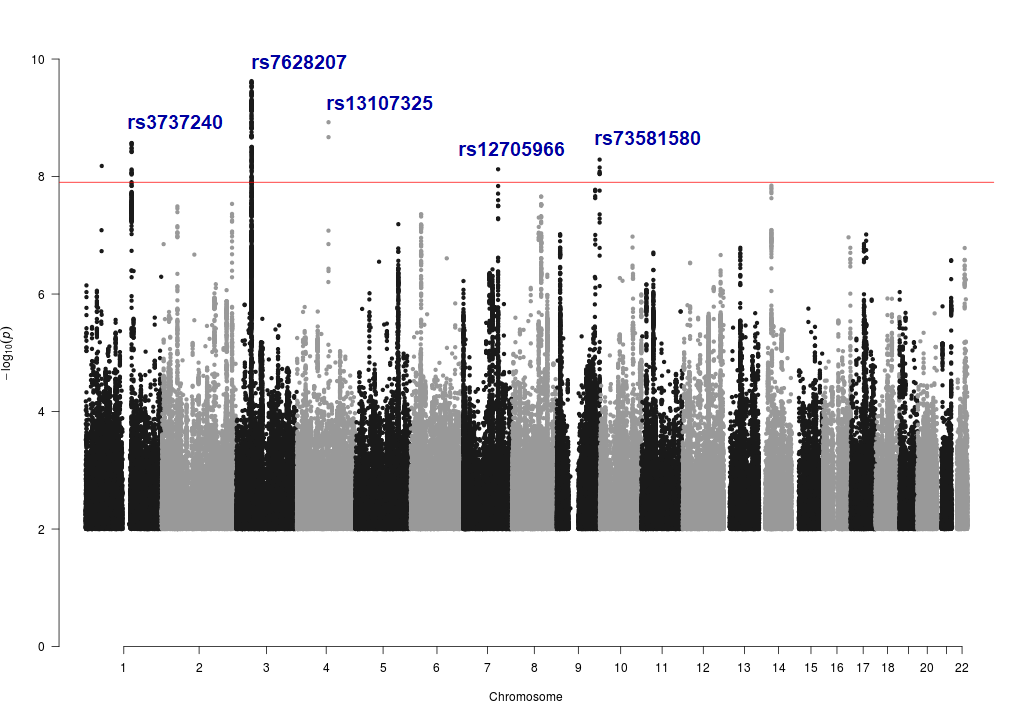

**GIP2**

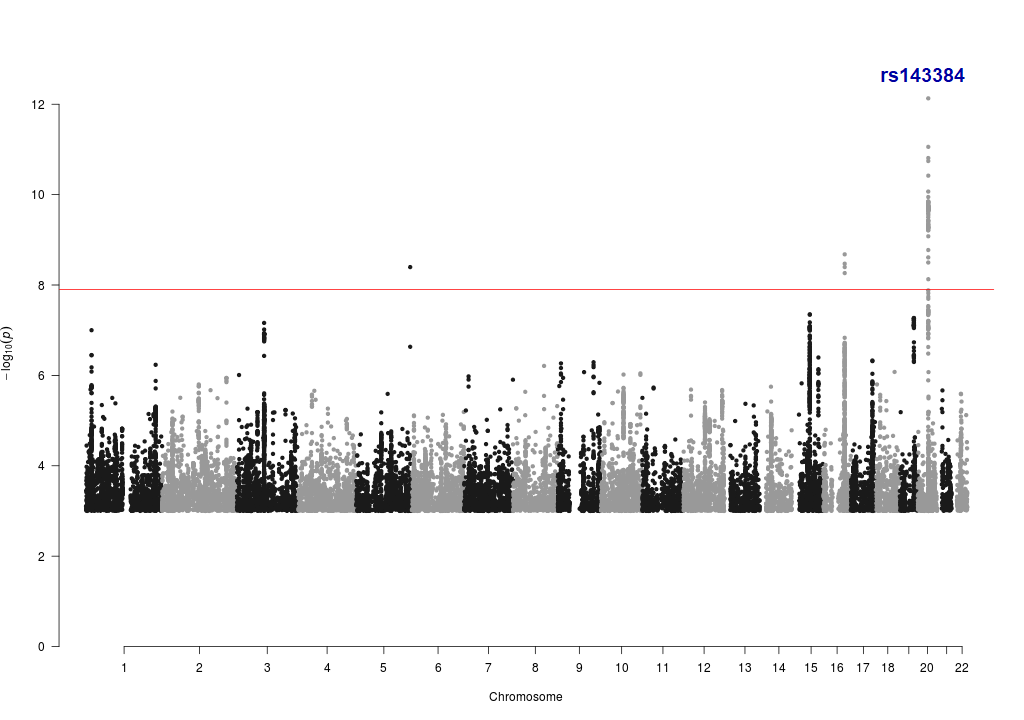

**GIP3**

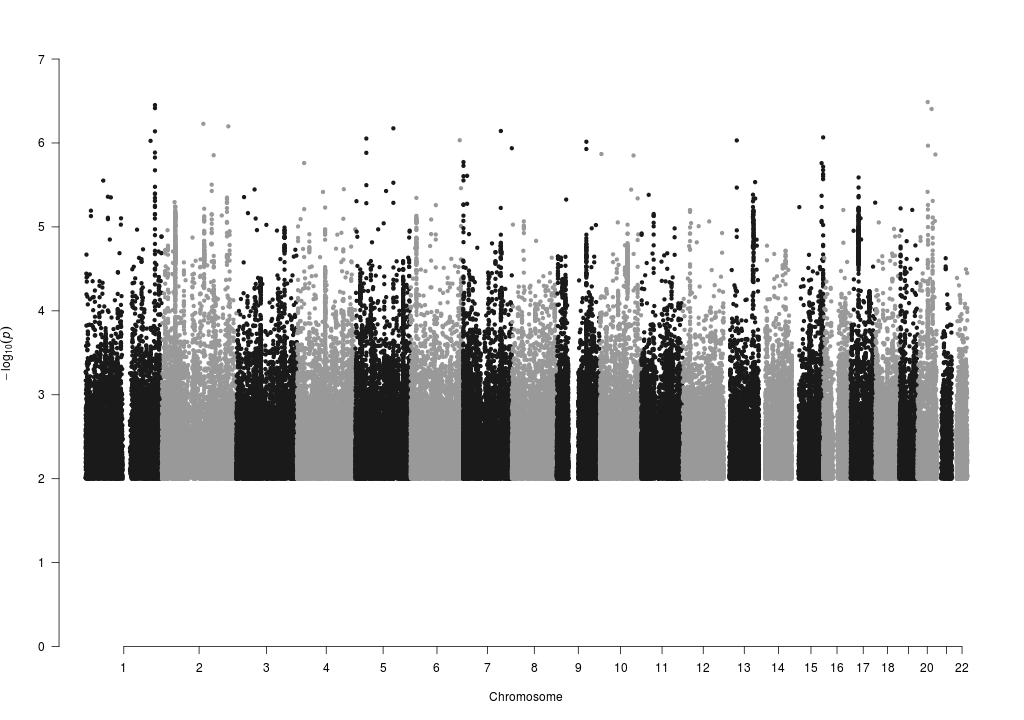

**GIP4**

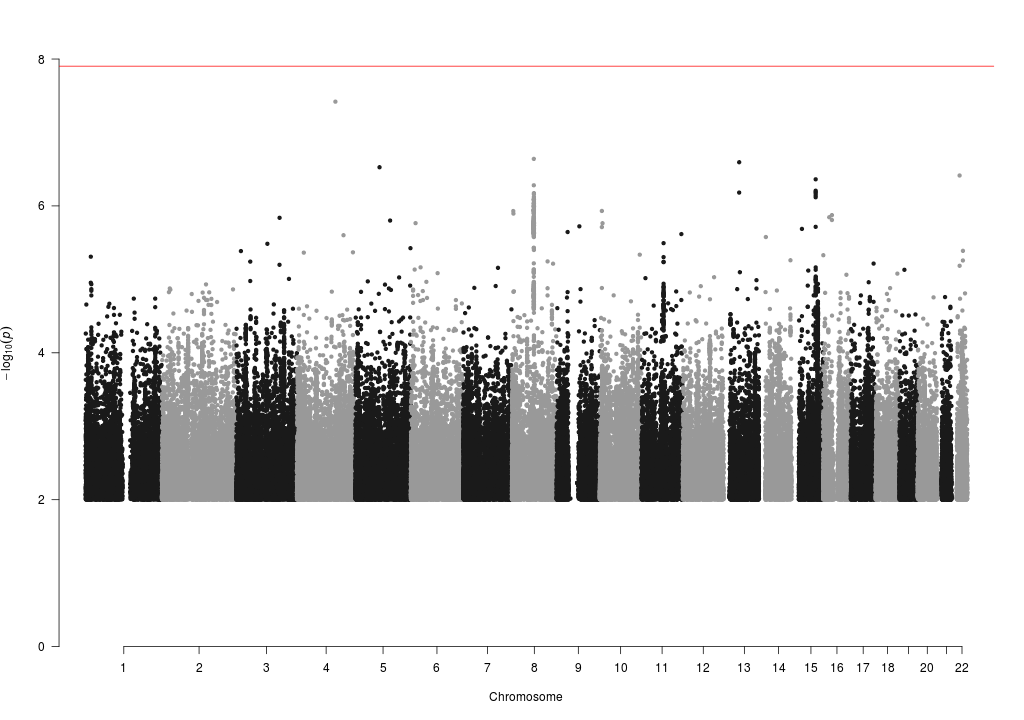

### Figure S2.

Quantile-quantile plots for observed vs. expected distribution of *P*-values for χ^2^ statistics.

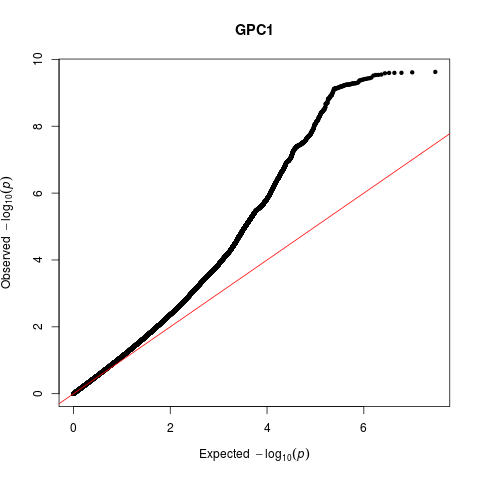

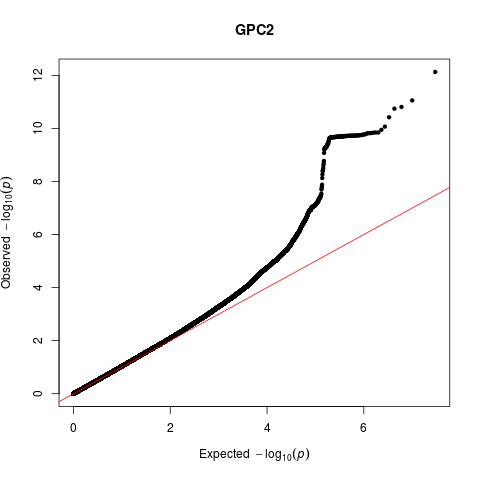

**GIP2**

**GIP1**

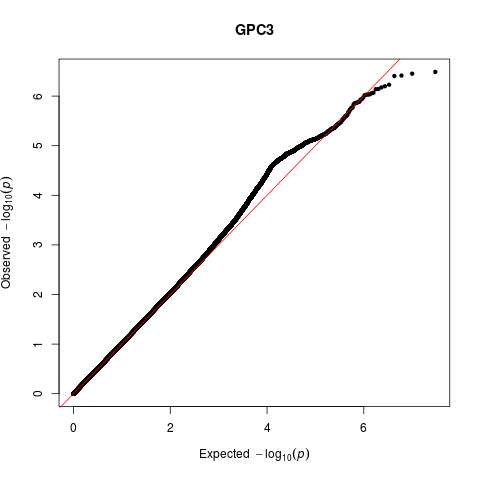

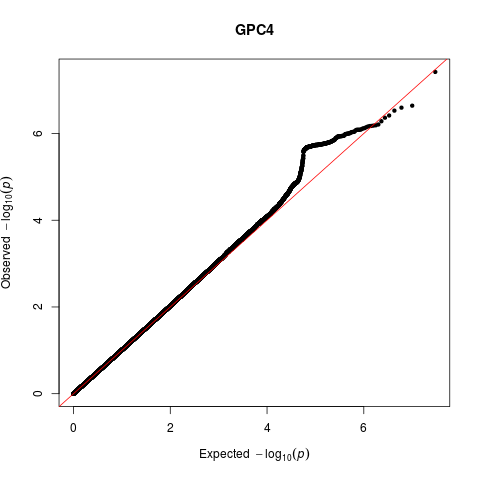

**GIP4**

**GIP3**

### Figure S3.

Regional association plots of –log_10_(*P*) for SNPs located at the distance of ≤ 250 kb from lead SNPs. Color of circles indicates the strength of linkage disequilibrium with the lead SNP based on the squared correlation coefficient (r^2^). Blue line indicates recombination rate (cM/Mb). Genes are indicated as blue bars under the plot.

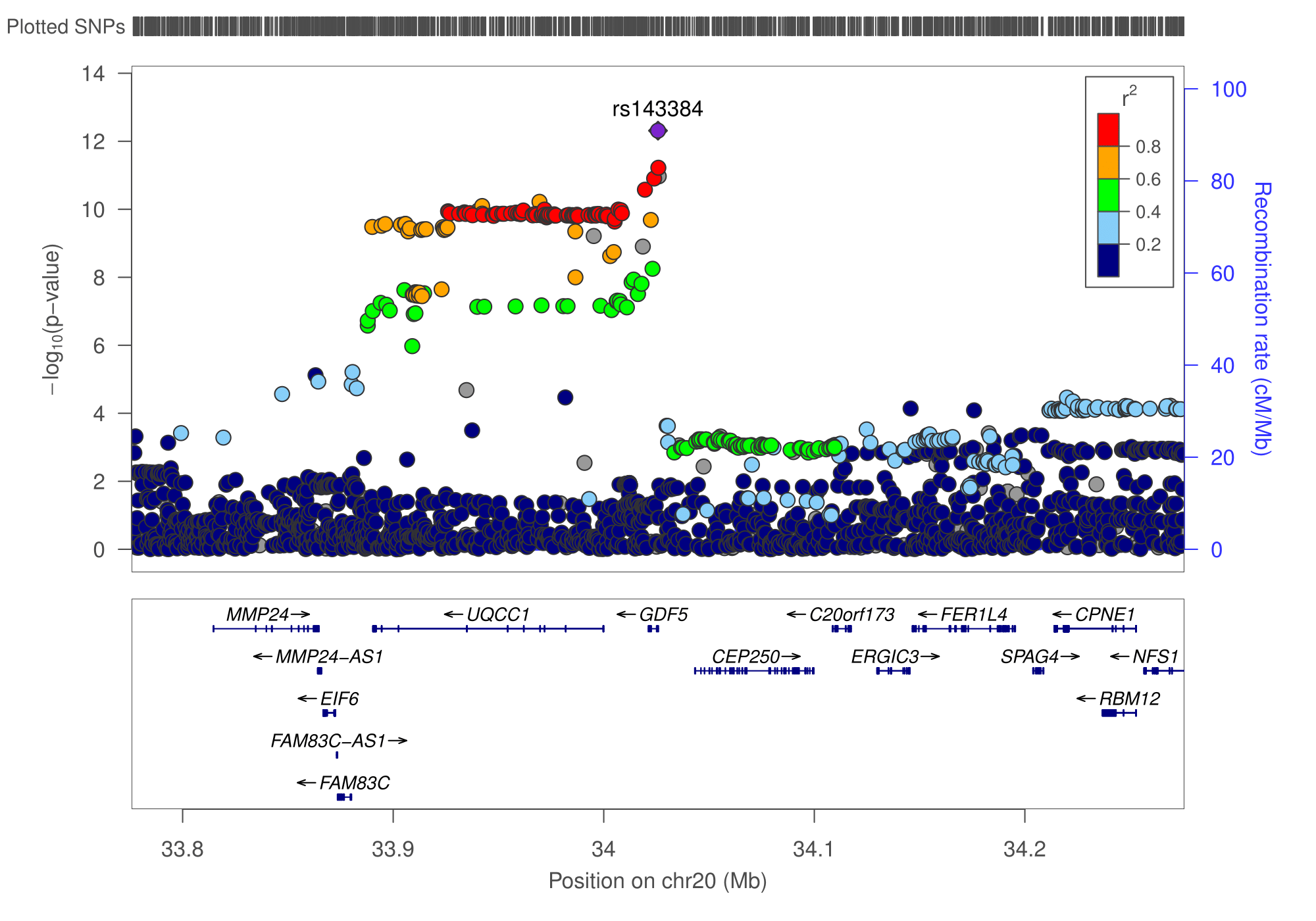

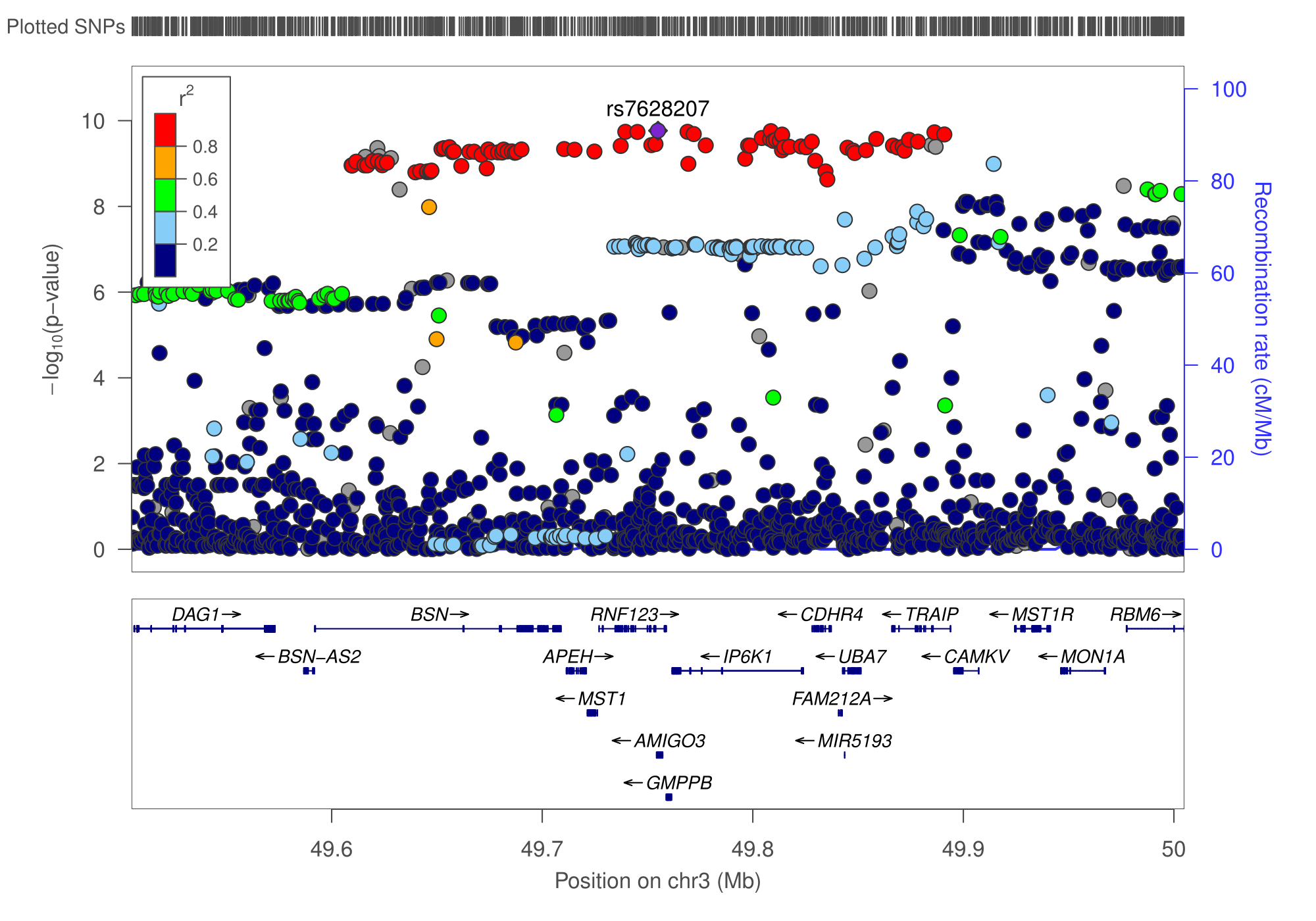

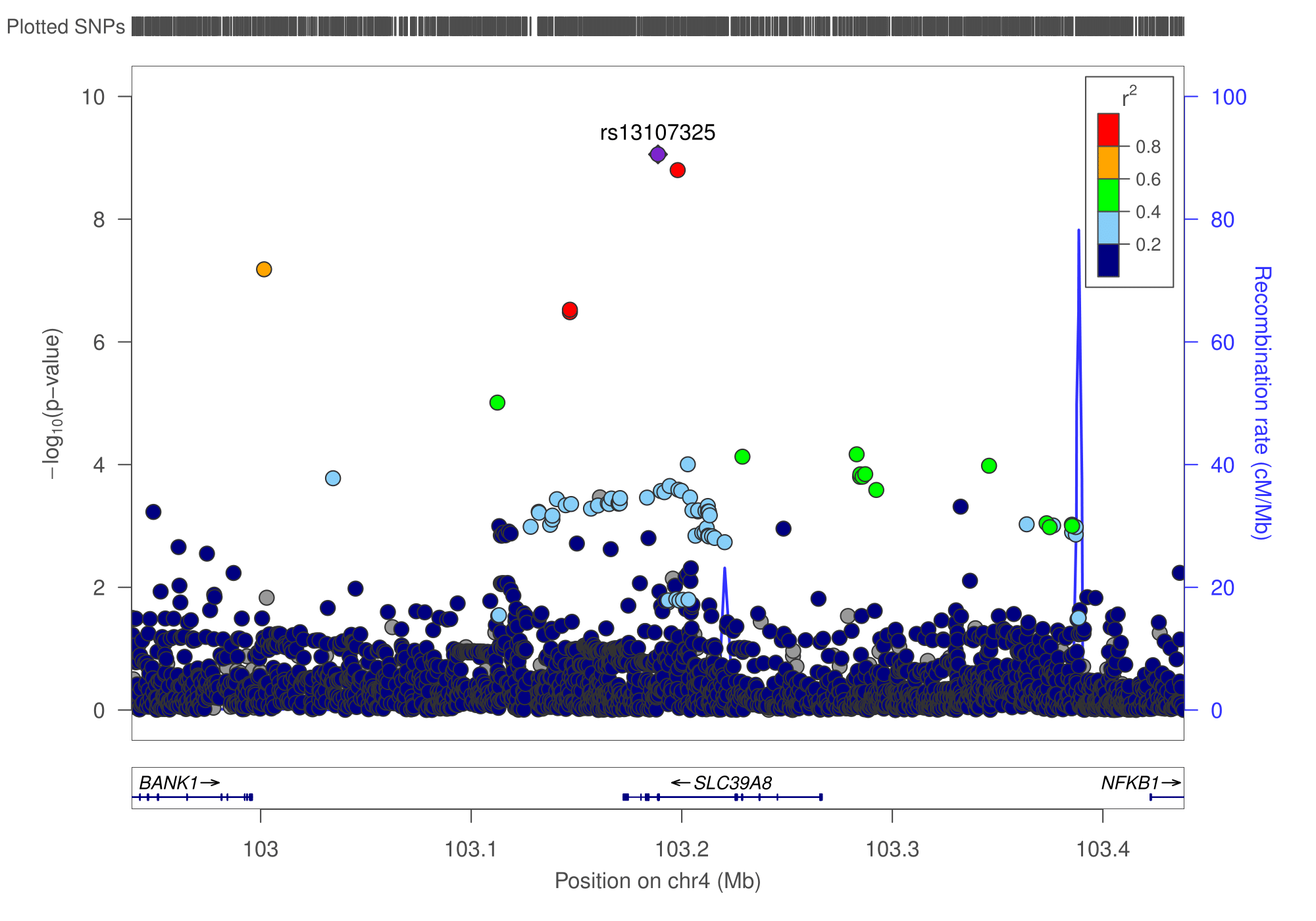

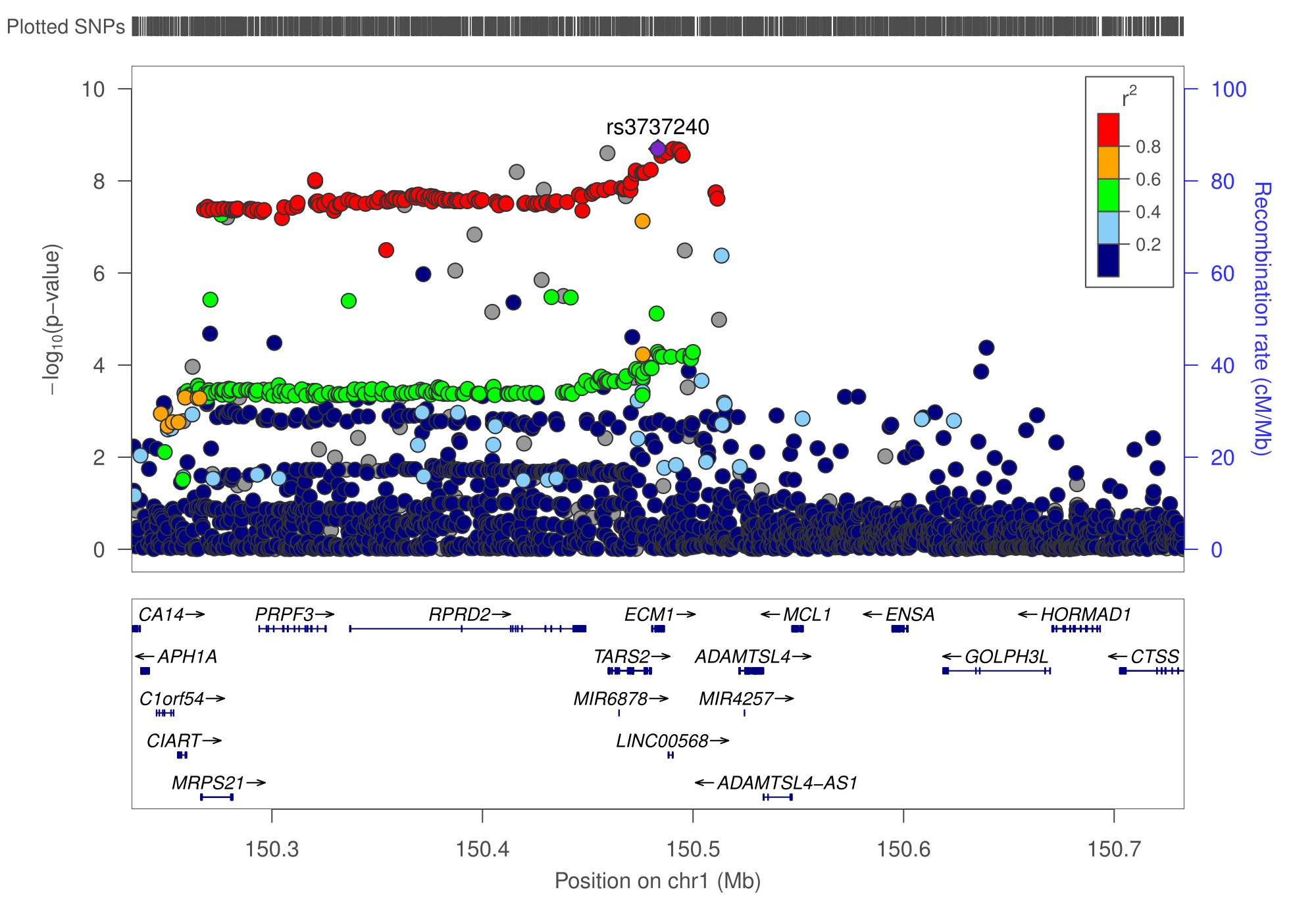

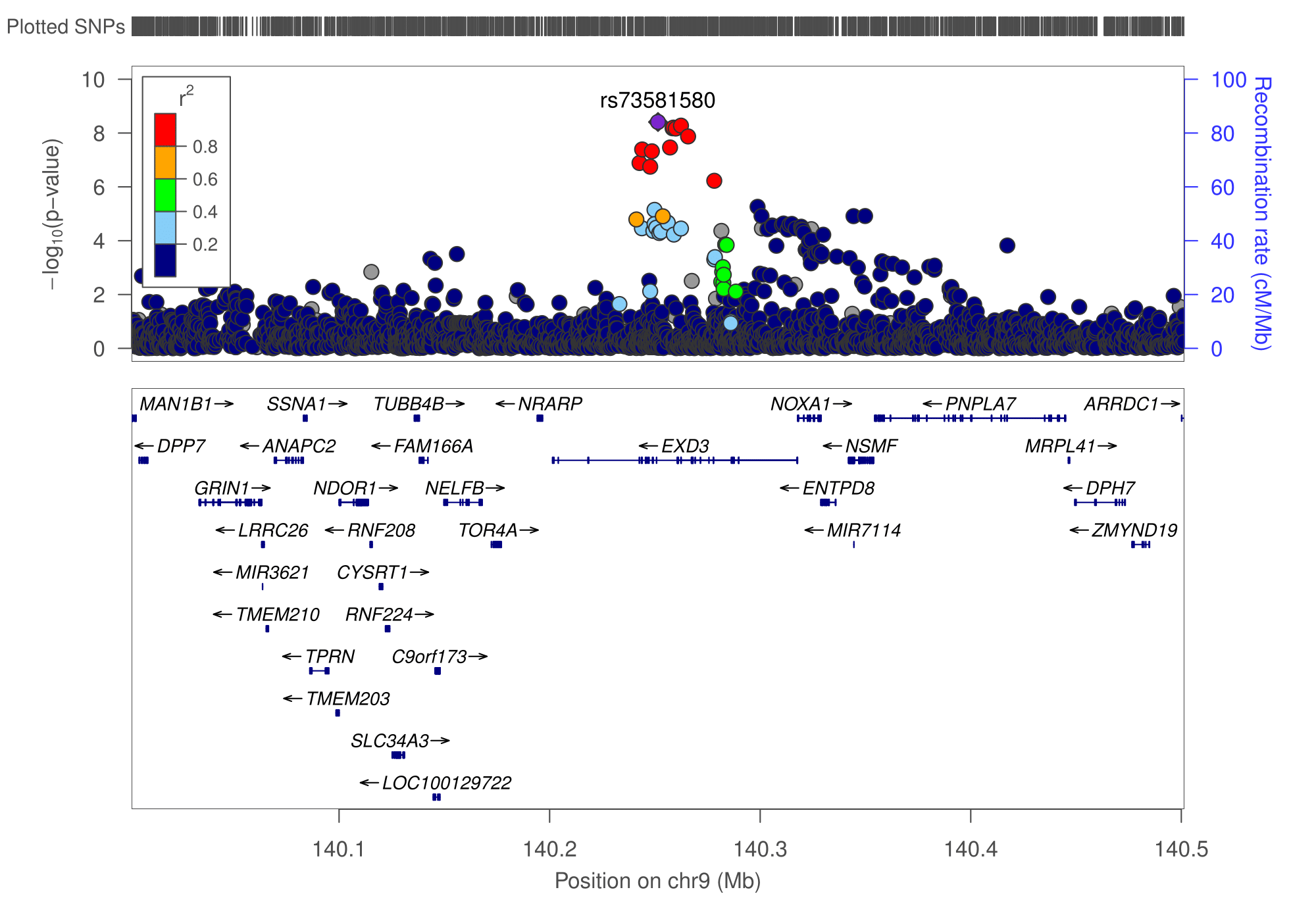

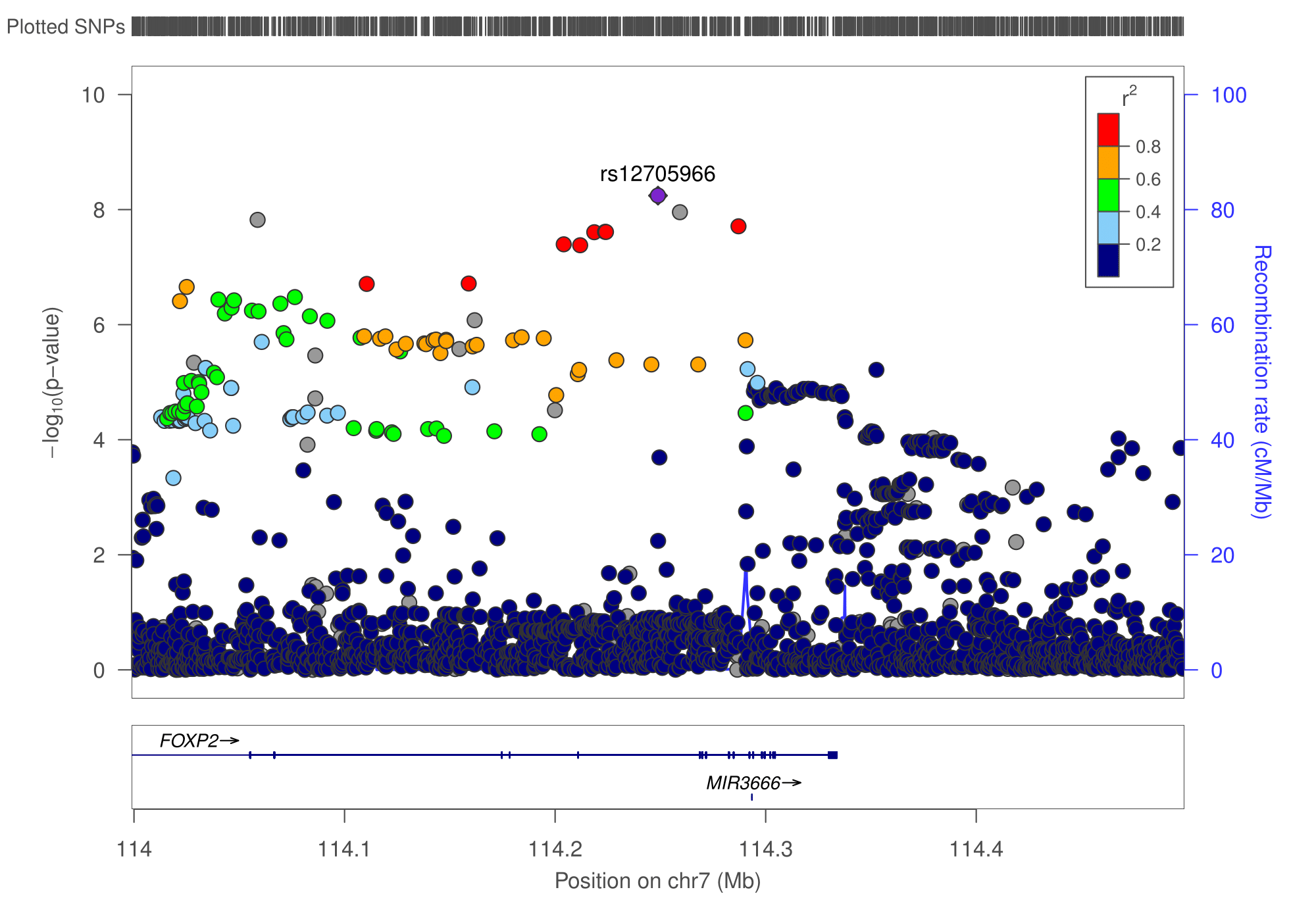

### Figure S4.

Results of tissue enrichment analysis for GIP1 performed using the FUMA platform (doi:10.1038/s41467-017-01261-5).

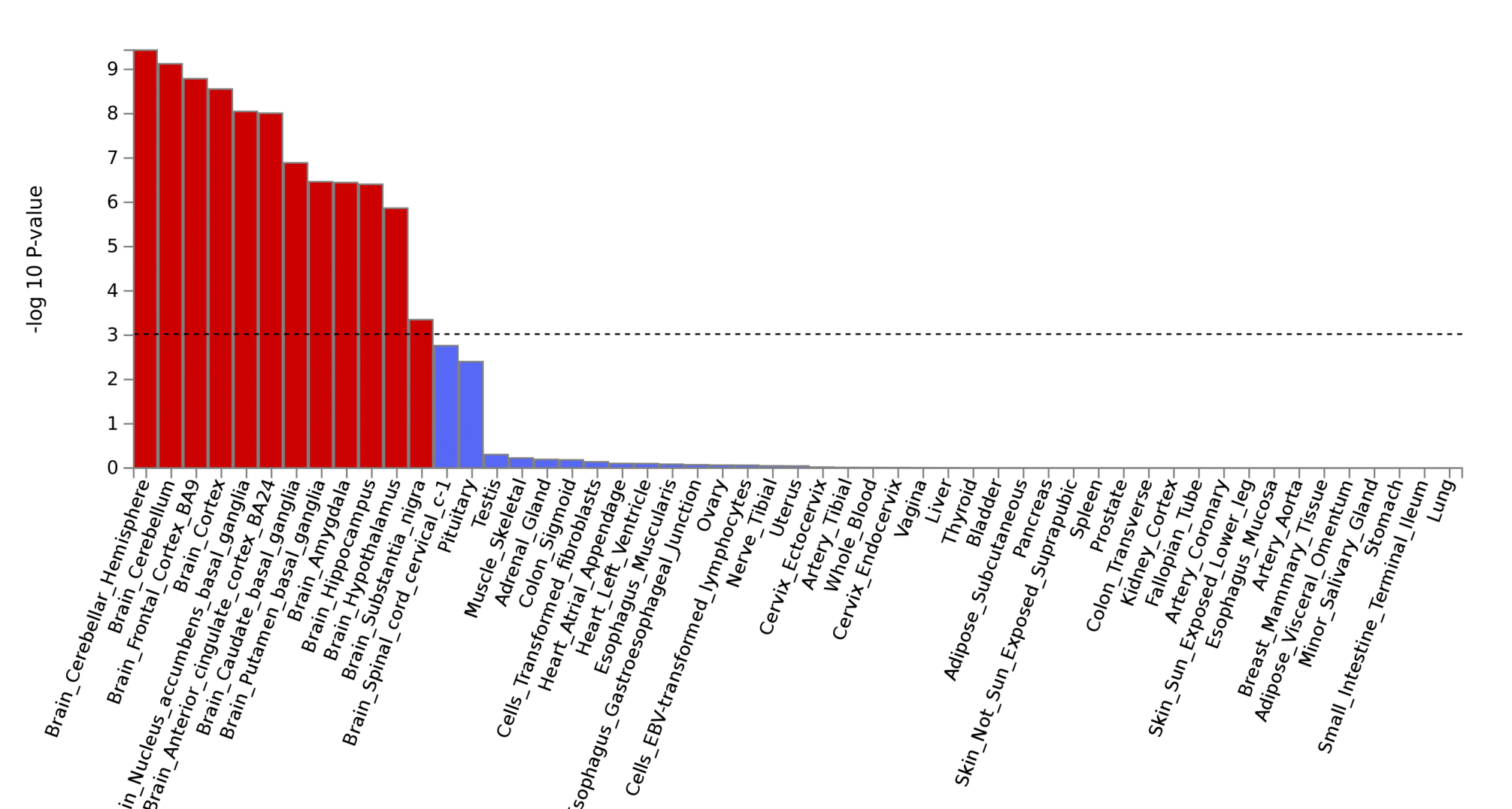

### Figure S5.

Matrix of genetic correlations between GIPs, chronic musculoskeletal pain traits and hospital-diagnosed osteoarthritis (the UK Biobank trait for which GWAS summary statistics was downloaded from the Michigan PheWeb database, <http://pheweb.sph.umich.edu/SAIGE-UKB/pheno/740>).

Color depicts the sign and absolute value of the genetic correlation coefficients (rg).

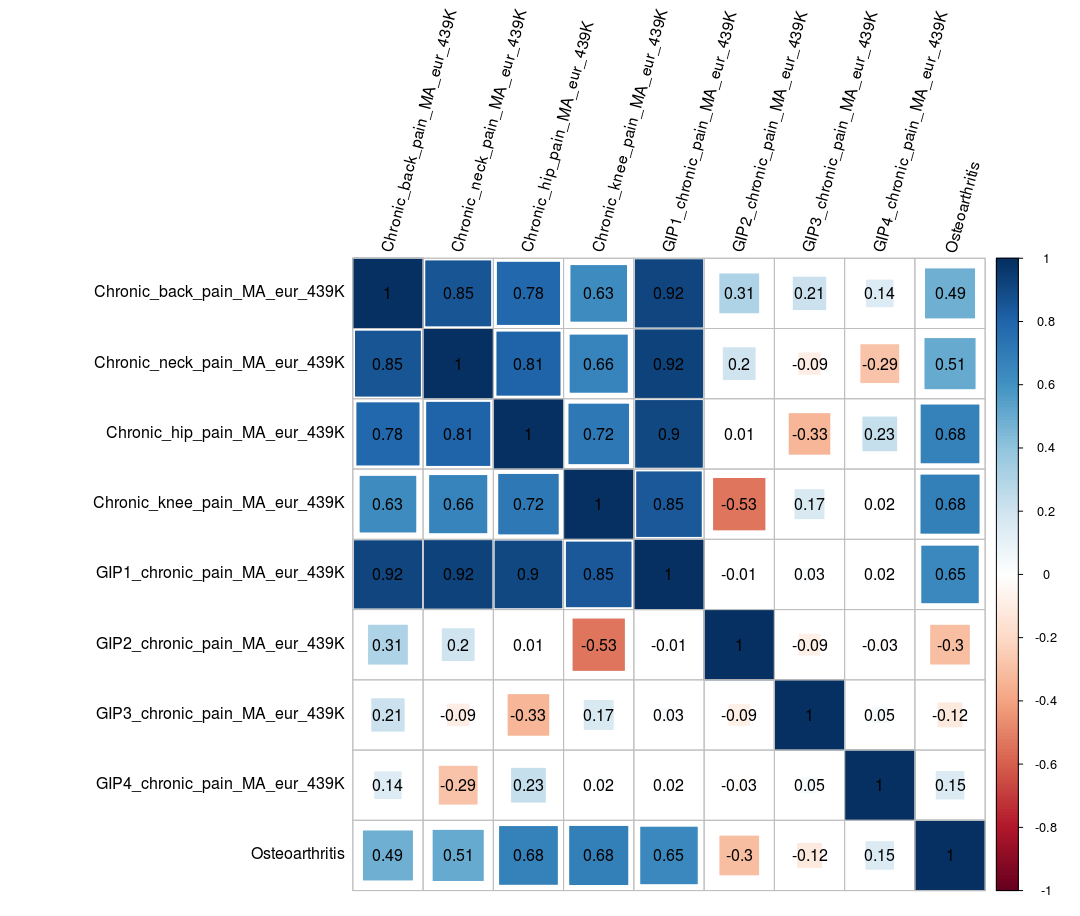
