## Supplementary Methods for "Analysis of genetically independent phenotypes identifies shared genetic factors associated with chronic musculoskeletal pain at different anatomic sites"

### Phenotype definition

Within the frame of the UK Biobank study, participants were asked to complete web-based questionnaires which included questions related to pain experience and duration.

We defined choric pain patients and corresponding controls based on their answers to “SY5” and “SY5B” questions^1^:

| **Q. No** | **Stem** | **Responses** | **Select from** |
| --- | --- | --- | --- |
| SY5 | In the last month have you experienced any of the following  that interfered with your usual activities?  (You can select more than one answer) | Select from  - 01 Headache  - 02 Facial pain  - 03 Neck or shoulder pain  - 04 Back pain  - 05 Stomach or abdominal pain  - 06 Hip pain  - 07 Knee pain  - 08 Pain all over the body  - NN None of the above  - DA Prefer not to answer | If any pain selected ask SY5B for that pain.  If none of the above or DA go to SY1 |
| SY5B | Have you had *** for more than 3 months?  **** for each pain selected in SY5 insert the response from SY5 with an ‘s’ following the word pain (eg, Have you had headaches for more than 3 months? Or Have you had neck or shoulder pains for more than 3 months? Or Have you had pains all over the body for more than 3 months?)* | Select from  - YE Yes  - NO No  - UN Do not know  - DA Prefer not to answer | Loop to go through SY5B for each pain selected (except ‘pain all over the body’ where only one selection allowed in SY5. Go to SY1 when all selections completed. |

- Individuals who preferred not to answer the SY5 question as well as individuals who selected “Pain all over the body” AND answered “Yes” to the SY5B question were excluded from the study.
- For each studied pain phenotype (back, neck/shoulder, knee, and hip), individuals who selected the corresponding pain in the SY5 section AND answered “Yes” to the SY5B question were defines as cases.
- Similarly, individuals who did not choose a specific pain in the SY5 section (either answering “None of the above” or selecting another pain) as well as individuals who selected the studied pain AND answered “No” to the SY5B question were defined as controls to each pain phenotype.

### GWAS-MAP platform

GWAS-MAP platform was originally created to study cardiovascular diseases. It integrates a database of summary-level GWAS results for different complex traits, including plasma/IgG N-glycome traits, levels of circulating metabolites, cytokines, growth factors, and other proteins. In the present study, we additionally added 18 chronic musculoskeletal pain-related traits and osteoarthritis . A brief description of data is given in Table SM1. All 2,262 traits are listed in Table S2.

SNPs from each GWAS were matched with polymorphisms from 1000 Genomes Project Phase 3 v5 reference panel. SNPs with conflicting data on rs id, position, and alleles were excluded. For SNPs that have passed this filtering, alleles were harmonized across all GWAS and sorted in a lexicographic order.

GWAS summary statistics is stored using the ClickHouse database management system (<https://clickhouse.yandex/>). For each trait, we created an annotation file that contains information about the study design and key characteristics of association analysis (name of the cohort, sample size, model of inheritance, trait transformations, reference population, etc.). Metadata is organized with the PostgreSQL database system (<https://www.postgresql.org/>).

Besides a GWAS database, our platform contains embedded software for LD Score regression^1^, 2-sample Mendelian randomization analysis^2,3^, and our implementation of SMR/HEIDI analysis^4^ (see below).

**Table SM1.** Data included in the GWAS-MAP database.

| **Dataset/source** | **Number of traits** | **Description** | | **Reference** |
| --- | --- | --- | --- | --- |
| The Neale Lab | 639 | Complex traits from the UK Biobank^*^ | | <http://www.nealelab.is/> |
| The Gene ATLAS | 34 | Complex traits from the UK Biobank^*^ | | <http://geneatlas.roslin.ed.ac.uk/> |
| “Metabolomics_NMR” | 123 | Circulating metabolites quantified with the NMR metabolomics platform (University Hospitals of Strasbourg, France) | | ^5^ |
| “Protein biomarkers_Olink” | 82 | Plasma proteins considered relevant to cardiovascular disease measured with the ProSeek CVD array I (Olink Biosciences, Sweden) | | ^6^ |
| “Proteomics_SOMAscan” | 1,124 | Blood circulating proteins measured with the SOMAscan platform (SomaLogic Inc., USA) | | ^7^ |
| “CAD_traits” | 8 | Coronary artery disease-related traits | Coronary artery disease | ^8^ |
|  |  |  | Coronary artery disease | ^9^ |
|  |  |  | Myocardial infarction | ^10^ |
|  |  |  | Fasting glucose | ^11^ |
|  |  |  | Cigarettes smoked per day | ^12^ |
|  |  |  | Body mass index | ^13^ |
|  |  |  | Waist-hip ratio | ^14^ |
|  |  |  | Educational attainment | ^15^ |
| “IBD” | 2 | Inflammatory bowel disease (Crohn's disease and ulcerative colitis) | | ^16^ |
| “Cytokines and growth factors” | 41 | Circulating cytokines and growth factors (measured in plasma or serum) | | ^17^ |
| “IgG glycome” | 77 | Plasma IgG N-glycome traits measured by UPLC | | <https://doi.org/10.7488/ds/2481> |
| “Plasma glycome” | 113 | Plasma N-glycome traits measured by UPLC | | <https://doi.org/10.5281/zenodo.1298406> |
| **Additionally added** | | | | |
| “Chronic pain” | 18 | Chronic pain-related traits | Chronic back, neck/shoulder, knee, hip pain (UK Biobank traits)^**^ (two datasets – discovery dataset and European ancestry meta-analysis – for each trait) | GWAS summary statistics was obtained in the present study |
|  |  |  | Four genetically independent phenotypes for back, neck/ shoulder, knee, and hip pain (two datasets – discovery dataset and European ancestry meta-analysis – for each phenotype) |  |
|  |  |  | Chronic stomach/abdominal pain; headache (UK Biobank traits)^**^ (European ancestry meta-analysis) |  |
| “Michigan PheWeb” | 1 | Osteoarthritis | The UK biobank trait. GWAS was performed using SAIGE method^18^, and the results were deposited in the Michigan PheWeb database (trait “740: Osteoarthrosis”). | <http://pheweb.sph.umich.edu/SAIGE-UKB/pheno/740>  ^18^ |

^*^ Binary traits with the number of cases or controls < 2000 were not included in the database.

- Data from the Neale Lab database were downloaded on December 15, 2017.
- Data from the Gene ATLAS were downloaded on December 8, 2017.

^**^ Genotyping and imputation data were obtained from the UK Biobank March 2018 data release under the project #18219 “Genetic and epidemiological analyses of low back pain”.

### Genetically independent phenotypes analysis – GIPA

To elucidate genetic component explaining most cases of four chronic musculoskeletal pain phenotypes, we proposed to use a modified **principal component analysis** (PCA) technique. PCA is a statistical procedure that uses an orthogonal transformation to convert a set of possibly correlated variables into a set of linearly uncorrelated variables called principal components (PCs). Each PC is a linear combination of the original variables. The first PC explains as much variability as possible (has the largest possible variance), and each succeeding component accounts for the largest proportion of the remaining variability under the constraint that it is orthogonal to the preceding components. The resulting vectors of orthogonal transformation coefficients (denoted as ***a_i_***) are an uncorrelated orthogonal basis set. PCA is sensitive to the relative scaling of the original variables. In the case of positive semidefinite covariance matrix of original variables (***∑***), ***a_i_*** are eigenvectors of **∑**. Each corresponding eigenvalue is proportional to the portion of the “variance explained” (more correctly of the sum of the squared distances of the points from their multidimensional mean) that is associated with each eigenvector.

To decompose the traits of interest into the genetically independent components, we proposed to use the matrix of genetic covariances **Ω** (instead of the matrix of phenotypic covariance used in conventional PCA for biological traits) for extraction of ***a_i_***. We termed the resulting principal components **“genetically independent phenotypes” (GIPs)**.

GIPs have several properties:

1. Each GIP is the specific linear combination of original traits. Thus, corresponding GWAS results can be obtained for each GIP, and each GIP can be analyzed as a separate trait using *in-silico* follow-up approaches.
2. GIPs are genetically independent from each other: pairwise genetic correlations between any GIPs are zero (albeit phenotypic correlations of original traits may not be equal to zero).
3. Confidence intervals (CI) for ***a_i_*** can be estimated using the standard errors of genetic covariance matrix estimation.

Technical details of the genetic principle component analysis approach are provided below.

Denote the following variables:

Ω – the matrix of genetic covariances (*m* x *m*, where *m* is the number of traits)

Ω_SE_ – the matrix of the standard errors of genetic covariances (*m* x *m*)

∑_ph_ – the matrix of phenotypic covariances (*m* x *m*); in the case of standardized traits it is equal to the matrix of phenotypic correlations

B – the matrix of effect sizes (*β*) for *m* phenotypes (*M* x *m*, where *M* is the number of SNPs in the analysis). *b_i_* is the *i*-th column of B.

SE – the matrix of standard errors of *β* for *m* phenotypes (*M* x *m*). *SE*_i_ is the *i*-th column of SE.

*varY*_i_ – the variance of the *i*-th trait. After standardization, *varY*_i_ = 1.

*SD_i_* – the standard deviation of *i*-th trait. ${SD}_{i}= \sqrt{{varY}_{i}}$

B_s_ – the matrix of standardized *β* for *m* phenotypes (*M* x *m*). b^s^_i_ is the *i*-th column of B_s_.

SE_s_ – the matrix of standardized standard errors for *m* phenotypes (*M* x *m*). *SE^s^_i_* is the *i*-th column of SE_s_.

A – the matrix of eigenvectors of Ω (*m* x *m*). Each column is *a_i_*, {*a_1_…a_m_*} – the vector of orthogonal transformation coefficients of *m* original traits into *m* GIPs {GIP_1_.. GIP_m_}

A_s_ – the matrix of scaled eigenvectors of Ω (*m* x *m*). Each column is *a^s^_i_*, {*a^s^_1_*…*a^s^_m_*} – the vector of orthogonal transformation coefficients of *m* original traits into *m* GIPs scaled to make the GIPs’ variance equal to 1.

*L* – the vector of eigenvalues {*l_1_…l_m_*}.

We performed the following procedure to calculate GIPs for four studied pain phenotypes:

1. Estimated Ω and Ω_SE_ using LDSC software (<https://github.com/bulik/ldsc/>).
2. Estimated *varY*_i_ and Pearson correlation matrix for four pain phenotypes.
3. Standardized GWAS summary statistics for four pain phenotypes ($\beta_{i}^{s}={\beta_{i}}/{{SD}_{i}}$ and ${SE}_{i}^{s}={{SE}_{i}}/{{SD}_{i}}$).
4. Checked whether all eigenvalues were positive for Ω.
5. Estimated eigenvalues (L) and the matrix of eigenvectors (A) of Ω.
6. If the coefficient of a given eigenvector for back pain was negative ($a_{i,back pain}$< 0), we changed the signs for all coefficients in its eigenvector ($a_{i}=-a_{i}$)
7. Estimated variance for GIP as ${var(GOP}_{i})=\sum\left[ (a_{i}\bigotimes a_{i})\circ\sum_{ph} \right]$, where $\bigotimes$ is an outer product.
8. Scaled coefficients for GIPs as $a_{i}^{s}={a_{i}}/{{SD(GOP}_{i})}$.
9. Estimated 95% CI for *a_i_* (see below)^*^.
10. Provided GWAS results for each GIP (see below)^**^.

*For estimation of 95% CI for GIPs, the Monte Carlo approach was used. We performed 1000 cycles of simulations. In each round, we simulated the noise component for matrix of genetic correlations Ω - the matrix Ω*^noise^* (*m* x *m*). Each element *i,j* (*i* > *j*) Ω*^noise^* is sampled from the normal distribution with zero mean and standard deviation equal to the *i,j* element of matrix of standard errors ($\Omega_{i,j}^{SE}$). The resulting covariance matrix was obtained as the sum of Ω and Ω*^noise^*. Than the standardized eigenvalues were calculated as described above formulating the empirical distribution of each element of A_s_ matrix. For each element of A_s_, 95% CI was obtained as an absolute difference between 0.975 and 0.025 quantiles divided by 2.

**Estimation of GWAS for GIPs was performed using the following procedure:

1. Effect sizes for *M* SNPs were calculated as $\beta_{{GOP}_{i}}=B_{s}\times a_{i}^{s}$, where $\times$ is an inner product.
2. Estimation of variance for GIPs was calculated as ${varGOP}_{i}=\sum\left[ (a_{i}\bigotimes a_{i})\circ\sum_{ph} \right]$. Phenotypic correlation matrix $\sum_{ph}$ was obtained from the specific studied population.
3. The standard errors of effect sizes for *M* SNPs were calculated as

${SE}_{{GPC}_{i}}=\sqrt{{varGOP}_{i}*\left( {{SE}_{1}^{s}}^{2}+\frac{{b_{1}^{s}}^{2}}{N} \right)- \frac{\beta_{{GOP}_{i}}^{2}}{N}}$, where $N$ is the sample size.

4. Scaling of effect sizes and standard errors were performed as described above.
5. The corresponding *P*-values were estimated using Wald test ($Z-score={\beta_{{GOP}_{i}}}/{{SE}_{{GOP}_{i}}}$).

- **The total genetic variance of m original traits explained by each GIP was calculated as** $\boldsymbol{R}_{\boldsymbol{GOP}_{\boldsymbol{i}}}^{\boldsymbol{2}}\boldsymbol{=}{\boldsymbol{l}_{\boldsymbol{i}}}/{\sum_{\boldsymbol{i}\boldsymbol{=1}}^{\boldsymbol{m}} \boldsymbol{l}_{\boldsymbol{i}}}$**.**
- The heritability of each GIP was calculated as $h_{{GOP}_{i}}^{2}=\frac{\sum\left[ (a_{i}\bigotimes a_{i})\circ\Omega\right]}{\sum\left[ (a_{i}\bigotimes a_{i})\circ\sum_{ph} \right]}$.
- Genetic correlations between GIPs and original traits as well as between each other were calculated as

$$\rho_{genetic}\left\{ c_{1} | c_{2} \right\}=\frac{\sum\left[ (c_{1}\bigotimes c_{2})\circ\Omega\right]}{\sqrt{\sum\left[ (c_{1}\bigotimes c_{1})\circ\Omega\right]\times\sum\left[ (c_{2}\bigotimes c_{2})\circ\Omega\right]}}$$

given that GIPs are linear combinations of original traits ($c_{1}$ and $c_{2}$, in case of ${GOP}_{i}$ $c_{j}=a_{i}^{s}$),

- Predicted phenotypic correlations between GIPs and original traits as well as between each other were estimated as

$$\rho_{phenotypic}\left\{ c_{1} | c_{2} \right\}=\frac{\sum\left[ (c_{1}\bigotimes c_{2})\circ\sum_{ph} \right]}{\sqrt{\sum\left[ (c_{1}\bigotimes c_{1})\circ\sum_{ph} \right]\times\sum\left[ (c_{2}\bigotimes c_{2})\circ\sum_{ph} \right]}}$$

given that GIPs are linear combinations of original traits ($c_{1}$ and $c_{2}$, in case of ${GOP}_{i}$ $c_{j}=a_{i}^{s}$),

- The contribution of each GIP into the genetic basis of original traits (the genetic variance explained by GIP) was estimated as squared genetic correlation coefficient of GIP with a given trait.

### Testing for pleiotropy using SMR/HEIDI approach

SMR/HEIDI analysis was conducted as described by Zhu et al.^1^ HEIDI statistics was calculated as $T_{HEIDI}= \sum_{i}^{m} z_{d(i)}^{2}$, where *m* is the number of SNPs selected for analysis, $z_{d\left( i \right)}= \frac{d_{i}}{{SE}_{{(d}_{i)}}}$ and $d_{i}= \beta_{{SMR}_{i}}- \beta_{SMR (lead SNP)}$.

SNP selection was performed as follows:

1) We defined a set of eligible markers within ±250 kb from the lead SNP in the primary GWAS, which had χ^2^ > 10 in the primary GWAS, and for which the results were reported in the secondary GWAS;
2) Made empty “target” and “rejected” SNP sets;
3) Selected SNP from the primary GWAS with the lowest *P*;
4) If this SNP had r^2^ > 0.9 with any SNP in the target SNP sethttps://www.cog-genomics.org/plink2, we added it to the “rejected” set. LD matrix (r^2^) was computed with PLINK 1.9 (<https://www.cog-genomics.org/plink2>) using 1000 Genomes data for 503 European individuals (<http://www.internationalgenome.org/data/>);
5) Otherwise, it was added to the “target” set;
6) Procedure was repeated from the step 3) until either eligible SNP set was exhausted, or the “target” set had 20 SNPs. If we could not select 3 or more SNPs, no test was performed.

When testing for pleiotropy with complex traits, we standardized all SNP effects (*β*) and standard errors (made the variances of the traits equal to 1):

$\breve{\beta}_{Y_{i}}={\beta_{Y_{i}}}/{{SD}_{Y_{i}}}$ and $\breve{SE}_{Y_{i}}={{SE}_{Y_{i}}}/{{SD}_{Y_{i}}}$, where $\breve{\beta}_{Y_{i}}$ and $\breve{SE}_{Y_{i}}$ are standardized betas and standard errors for the trait $Y_{i}$; $\beta_{Y_{i}}$ and ${SE}_{Y_{i}}$ are original betas and standard errors for the trait $Y_{i}$; ${SD}_{Y_{i}}$is a square root of estimated variance of the trait $Y_{i}$. Analysis was conducted using Python 3.5 as the main programming language.
