## Supplementary Tables for "Analysis of genetically independent phenotypes identifies shared genetic factors associated with chronic musculoskeletal pain at different anatomic sites": Table S1_study cohort.docx

**Table S1.** Descriptive characteristics of the study cohorts.

|  | Prevalence | Sample size | Age (mean ± SD), *years* | BMI (mean ± SD), *kg/m^2^* | Women, *%* |
| --- | --- | --- | --- | --- | --- |
| **Discovery cohort* (N = 265,000)** | | | | | |
| Chronic back pain | 17.9% | Cases (N = 47,507) | 57.65 (7.99) | 28.33 (5.18) | 53.88 |
|  |  | Controls (N = 217,493) | 57.26 (8.03) | 27.15 (4.61) | 54.32 |
| Chronic neck pain | 16.3% | Cases (N = 43,287) | 57.73 (7.79) | 27.90 (5.02) | 53.84 |
|  |  | Controls (N = 221,713) | 57.25 (8.07) | 27.25 (4.68) | 54.32 |
| Chronic hip pain | 9.2% | Cases (N = 24,300) | 59.15 (7.44) | 28.91 (5.40) | 54.35 |
|  |  | Controls (N = 240,700) | 57.15 (8.06) | 27.20 (4.64) | 54.23 |
| Chronic knee pain | 17.5% | Cases (N = 46,292) | 58.61 (7.59) | 29.18 (5.37) | 54.12 |
|  |  | Controls (N = 218,708) | 57.06 (8.09) | 26.97 (4.50) | 54.27 |
| **Replication cohort (N = 191,580)** | | | | | |
| ***African ancestry* (N = 7,541)** | | | | | |
| Chronic back pain | 21.0% | Cases (N = 1,586) | 53.77 (8.24) | 30.62 (5.79) | 54.50 |
|  |  | Controls (N = 5,955) | 52.04 (8.00) | 29.27 (5.13) | 54.19 |
| Chronic neck pain | 16.1% | Cases (N = 1,217) | 54.38 (7.98) | 30.06 (5.52) | 54.353 |
|  |  | Controls (N = 6,324) | 52.02 (8.04) | 29.45 (5.25) | 54.24 |
| Chronic hip pain | 8.5% | Cases (N = 641) | 55.00 (7.91) | 31.30 (6.14) | 54.37 |
|  |  | Controls (N = 6,900) | 52.16 (8.05) | 29.39 (5.19) | 54.25 |
| Chronic knee pain | 20.4% | Cases (N = 1,539) | 54.67 (8.30) | 31.64 (6.11) | 54.49 |
|  |  | Controls (N = 6,002) | 51.82 (7.92) | 29.01 (4.93) | 54.20 |
| ***European ancestry* (N = 174,831)** | | | | | |
| Chronic back pain | 18.0% | Cases (N = 31,428) | 57.62 (7.96) | 28.36 (5.22) | 54.05 |
|  |  | Controls (N = 143,403) | 57.26 (8.02) | 27.14 (4.58) | 54.28 |
| Chronic neck pain | 16.3% | Cases (N = 28,482) | 57.82 (7.76) | 27.92 (5.02) | 54.27 |
|  |  | Controls (N = 146,349) | 57.23 (8.06) | 27.25 (4.66) | 54.24 |
| Chronic hip pain | 9.2% | Cases (N = 16,022) | 59.26 (7.40) | 28.86 (5.41) | 54.61 |
|  |  | Controls (N = 158,809) | 57.13 (8.05) | 27.21 (4.63) | 54.20 |
| Chronic knee pain | 17.3% | Cases (N = 30,173) | 58.71 (7.54) | 29.24 (5.41) | 54.27 |
|  |  | Controls (N = 144,658) | 57.04 (8.08) | 26.97 (4.47) | 54.23 |
| ***South Asian ancestry*** (N = 9,208)** | | | | | |
| Chronic back pain | 21.6% | Cases (N = 1,993) | 54.66 (8.51) | 27.76 (4.58) | 54.29 |
|  |  | Controls (N = 7,215) | 53.87 (8.47) | 26.92 (4.23) | 54.22 |
| Chronic neck pain | 20.2% | Cases (N = 1,864) | 54.65 (8.24) | 27.43 (4.56) | 54.31 |
|  |  | Controls (N = 7,344) | 53.88 (8.53) | 27.01 (4.25) | 54.22 |
| Chronic hip pain | 6.6% | Cases (N = 610) | 56.61 (8.21) | 28.30 (4.90) | 54.07 |
|  |  | Controls (N = 8,598) | 53.86 (8.47) | 27.01 (4.26) | 54.25 |
| Chronic knee pain | 20.1% | Cases (N = 1,850) | 55.97 (8.23) | 28.52 (4.86) | 54.10 |
|  |  | Controls (N = 7,358) | 53.55 (8.47) | 26.74 (4.09) | 54.27 |

*Discovery cohort comprised only individuals of European ancestry

**Indian, Pakistani, and Bangladeshi
