## Supplementary Tables for "Analysis of genetically independent phenotypes identifies shared genetic factors associated with chronic musculoskeletal pain at different anatomic sites": Table S6_literature-based prioritization.docx

**Table S6.** Gene prioritization based on a literature review. Protein names are indicated in italics.

| **Lead SNP** | **Candidate gene** | **OMIM^*^ code** | **Nearest gene?** | **Functional effects of the encoded proteins** |
| --- | --- | --- | --- | --- |
| rs143384 | *GDF5*  (*CDMP1*) | 601146 | **YES**; utr variant 5 prime | *Growth differentiation factor 5*  Regulates the development of numerous tissue and cell types, including cartilage, joints, brown fat, teeth, and the growth of neuronal axons and dendrites. GDF5 plays an important role in knee morphology [1]. *GDF5* gene has multiple different control sequences that show striking specificity for joints in the head, vertebral column, shoulder, elbow, wrist, hip, knee, and digits [2].  Mutations in the *GDF5* genes underlie rare skeletal disorders including Chondrodysplasia Grebe type [3], Acromesomelic chondrodysplasia, Hunter-Thompson type [4], and others. |
|  | *MMP24* | 604871 | NO; 160 kb from lead SNP | *Matrix metallopeptidase 24*  MMP-24 is involved in the proteolytic degradation of extracellular matrix in normal physiological processes as well as in disease such as arthritis and metastasis.  Evidence from mice studies:   - MMP-24 (MT5-MMP) is expressed in differentiated neurons and regulates axonal growth [5] - MMP-24 is an essential mediator of peripheral thermal nociception and inflammatory hyperalgesia [6] - Expression of MMP-24 immediately and gradually increased in the spinal cord following peripheral partial sciatic nerve ligation [7] - MMP-24 is essential for the development of mechanical allodynia and plays an important role in neuronal plasticity [8] |
| rs7628207 | *AMIGO3* | 615691 | **YES**; intron variant | *Adhesion molecule with Ig like domain 3*  AMIGO3 participates in the NgR1-p75/TROY receptor complex (substitutes for LINGO-1 protein). NgR1-p75/TROY-AMIGO3 mediates myelin-induced inhibition of axon growth in the acute phase of adult central nervous system injury [9].  Suppression of AMIGO3 disinhibits the growth of axotomized dorsal root ganglion neurons and enables neurotrophin-3 to stimulate the regeneration of spinal cord dorsal column axons [10]. |
|  | *BSN* | 604020 | NO; 46 kb from lead SNP | *Bassoon presynaptic cytomatrix protein*  BSN is thought to be a scaffolding protein involved in organizing the presynaptic cytoskeleton. *BSN* gene is expressed primarily in neurons in the brain.  In mutant mice, loss of BSN causes a reduction in normal synaptic transmission [11]. |

| rs13107325 | *SLC39A8*  *(ZIP8)* | 608732 | **YES**; missense | *Solute carrier family 39 member 8*  Transmembrane protein that acts as a transporter of several divalent cations, including manganese (Mn^2+^), zinc (Zn^2+^), cadmium (Cd^2+^), and iron (Fe^2+^) across the plasma membrane.  SLC39A8 was proposed to be involved in osteoarthritis-related cartilage degradation through modulation of Zn^2+^ concentration, which is required for catalytic activity of MMP-13 [12]. SLC39A8 (ZIP8) was upregulated in osteoarthritis cartilage of humans and mice. Ectopic expression of SLC39A8 in mouse cartilage tissue caused osteoarthritis cartilage destruction, whereas Zip8 knockout suppressed surgically induced osteoarthritis pathogenesis [13]. Down-regulating of SLC39A8 was shown to reduce cartilage destruction, that could be gained through targeting SLC39A8 by microRNA-488 [12].  In a zebrafish model, disruptive *slc39a8* gene mutation caused spinal abnormalities including thoracic spinal curvature and caudal vertebral fusions, impaired growth, and decreased motor activity [14]. |
| --- | --- | --- | --- | --- |
| rs3737240 | *MIR6878* | N/A | NO; 18 kb from lead SNP | MicroRNA 6878 was found to be differentially expressed in synovial fluid of male patients with osteoarthritis [15]. |
|  | *ECM1* | 602201 | **YES**; missense | *Extracellular matrix protein 1*  ECM1 is involved in endochondral bone formation and cartilage development (is a negative regulator of bone mineralization and chondrogenesis) [16-18], promotes angiogenesis [19], inhibits MMP9 proteolytic activity [20], interacts with many extracellular and structural proteins and contributes to the maintenance of skin integrity [21], is a potential activator of NF-kB signaling [22]. |
|  | *CTSS* | 116845 | NO; 219 kb from lead SNP | *Cathepsin S*  Lysosomal cysteine proteinase that can remodel extracellular matrix in various tissues.  Cathepsin S is critical for the maintenance of neuropathic pain and spinal microglia activation [23-26].  Cathepsin S is responsible for Th1 cell-dependent transition of nerve injury-induced acute pain to a chronic pain state [27]. |
| rs73581580 | *MIR7114* | N/A | NO; 93 kb from lead SNP | MicroRNA 7114 was found to be differentially expressed in T-cells from patients with ankylosing spondylitis [28]. |
|  | *NSMF (NELF)* | 608137 | NO; 91 kb from lead SNP | *NMDA receptor synaptonuclear signaling and neuronal migration factor*  NSMF was proposed to be involved in guidance of olfactory axon projections and migration of luteinizing hormone-releasing hormone neurons [29].  Nuclear import of NSMF (Jacob protein) is important for dendrite development in the hippocampus [30]. |
|  | *NOXA1* | 611255 | NO; 66 kb from lead SNP | *NADPH oxidase activator 1*  NOXA1 (member of the Nox family) activates NADPH oxidases, which catalyze a reaction generating reactive oxygen species.  Animal studies suggest that Nox enzymes contribute to signaling pathways involved in chronic inflammatory and/or neuropathic pain states [31]. |
|  | *GRIN1* | 138249 | NO; 218 kb from lead SNP | *Glutamate ionotropic receptor NMDA type subunit 1*  Critical subunit of N-methyl-D-aspartate receptors. These subunits play an important role in the plasticity of synapses, which is believed to underlie memory and learning [32].  A target for analgesic drugs [33]. |
| rs12705966 | *FOXP2* | 605317 | **YES**; intron variant | *Forkhead box P2*  FOXP2 is required for proper development of speech and language regions of the brain during embryogenesis. Mutations in FOXP2 cause developmental speech and language disorders in humans [34, 35]. |

^*^Online Mendelian Inheritance in Man database (<https://www.omim.org/>)
